# Symmetry reduction in a hyperpolarization-activated homotetrameric ion channel

**DOI:** 10.1101/2021.09.20.461140

**Authors:** Miles Sasha Dickinson, Meghna Gupta, Sergei Pourmal, Maxine Bi, Robert M. Stroud

**Author notes:** These authors contributed equally and are listed alphabetically (Miles Sasha Dickinson, Meghna Gupta, Sergei Pourmal).

## Abstract

Plants obtain nutrients from the soil via transmembrane transporters and channels in their root hairs, from which ions radially transport in towards the xylem for distribution across the plant body. We determined structures of the hyperpolarization-activated channel AKT1 from *Arabidopsis thaliana*, which mediates K^+^ uptake from the soil into plant roots. These structures of AtAKT1 embedded in lipid nanodiscs show that the channel undergoes a reduction of C4 to C2 symmetry, possibly to regulate its electrical activation.

## Main Text

Ion channels in plants perform a number of critical physiological tasks including the uptake of nutrients from the soil for plant development and viability, regulation of turgor pressure to control mechanical processes, and action potential-like electrical signaling ^1–3^. Among the most important nutrients for plants is K^+^, for which there exist numerous bona-fide channel genes in the model organism *Arabidopsis thaliana* ^1,3^ including 6-transmembrane helix (TM) *Shaker*-like channels, 12-TM two-pore channels, 4-TM twin pore channels, and 2-TM K_ir_ channels ^1,2^. The AKT1 channel is necessary for radial K^+^ transport, where it mediates the translocation of K^+^ from the rhizosphere into the root hair, from which the ion can diffuse through an outwardly-rectifying Stelar K+ outward rectifying (SKOR) channel into the xylem for transport across the plant shoot ^1,4,5^. The *Arabidopsis thaliana akt1* gene - first discovered by phenotypic rescue of K^+^ uptake-deficient yeast ^6^ - encodes a 96,990 Dalton α subunit that assembles into a homo-tetrameric channel. The protein contains four distinct domains: 1) a Shaker-like transmembrane cassette consisting of TM helices S1-S4 (constituting the voltage sensing domain) and S5-S6 (the pore domain); 2) a cyclic nucleotide binding homology domain (CNBHD); 3) an Ankyrin repeat domain; and 4) a C-terminal K_HA_ interaction domain.

In contrast to other putative cyclic nucleotide binding channels, AKT1 is unlikely to be regulated by cyclic nucleotide monophosphates (cNMPs) as it lacks critical residues for ligand binding ^7^ (**S4a**). Instead, channel gating is voltage dependent and modulated by a phosphorylation ‘switch’ ^1,3,8–10^. Yeast two-hybrid experiments show that the AKT1 Ankyrin repeat domain associates with a CIPK23 kinase / calcineurin B-like (CBL) calcium sensor complex, and patch clamp in *X. laevis* oocytes confirms that phosphorylation by this complex converts a nearly silent channel to a strongly hyperpolarization-activated channel with an activation threshold ~ −50 mV ^8^. Conversely, AKT1 can be inhibited by direct interaction with CBL10, or dephosphorylation by PP2C phosphatases ^8,11^. Phosphorylation couples channel activity, and thus K^+^ uptake, to intricate Ca^2+^ signaling networks in order for the plant to adapt to a range of soil conditions. Additionally, AKT1 is known to hetero-oligomerize with the pseudo-channel KC1, negatively shifting its V_1/2_ by ~ 60 mV ^11^. These modifications and interactions are critical for plant viability: activating modifications are necessary to facilitate the influx of K^+^ against large concentration gradients when [K^+^]_soil_ is low, and inhibition is needed to prevent K^+^ leakage out of the roots when E_K_ is below V_m_.

Here we present structures of the AKT1 channel from *A. thaliana*, determined by single particle cryogenic electron microscopy (cryoEM) (**1a,b**). We reconstituted AKT1 into soy polar lipid nanodiscs and determined a consensus structure to an average resolution of 2.7 Å. In addition, we recovered two other medium-resolution reconstructions of alternate flexible conformations, into which we docked consensus domains. Throughout our three structures, we observe a reduction of symmetry from the predicted C4 to C2, occurring in regions of the protein with known functional significance, namely the C-linker and CNBHDs. These structures reveal that the CNBHDs are highly mobile domains that flex about the channel axis, and that the C-linker alternates between two distinct conformations. Although the electrophysiological implications of these observations are yet to be determined, we suspect that similar processes may occur in other channels from the CNG and hyperpolarization-activated families.

**Figure 1.**
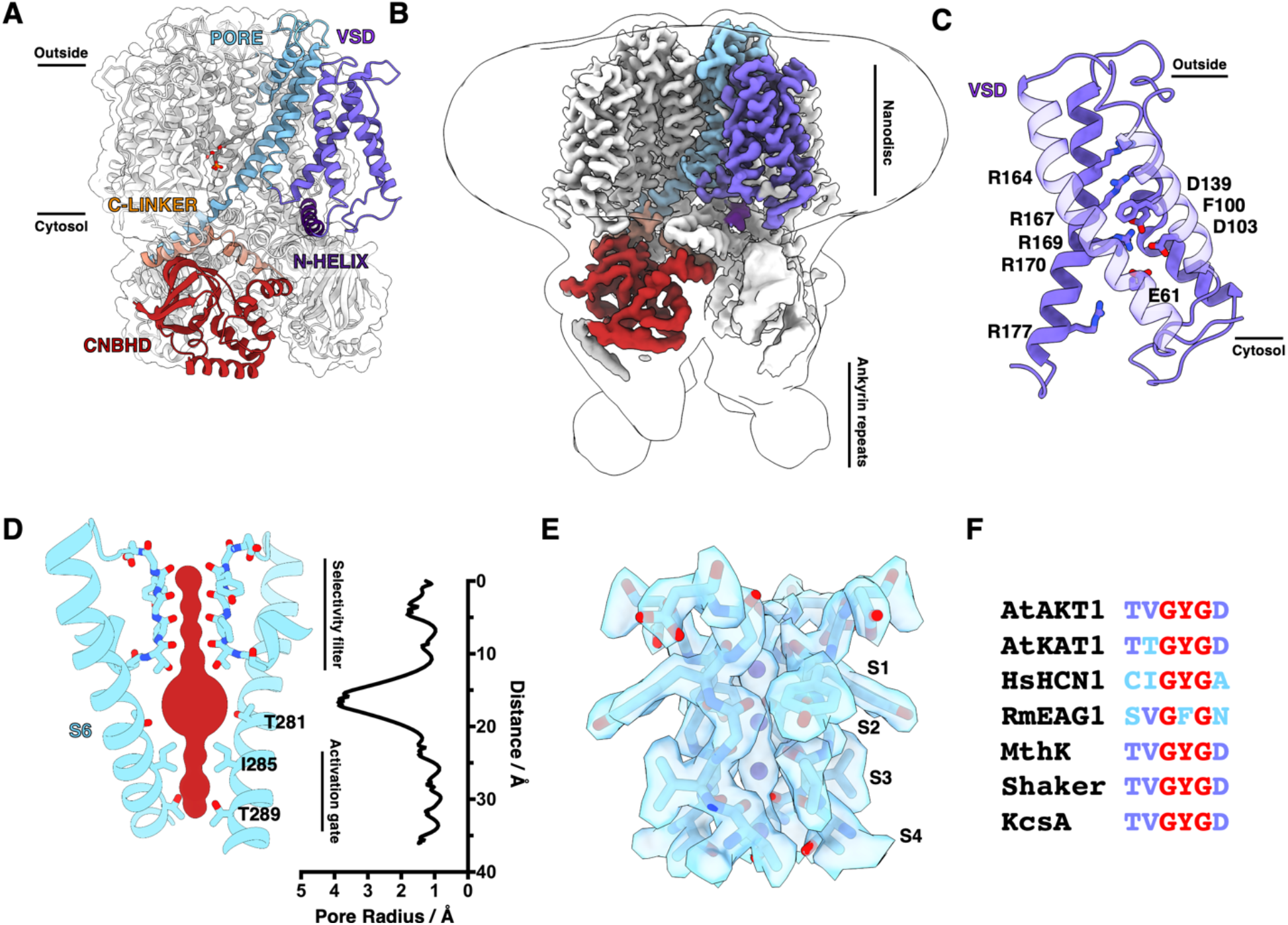
CryoEM structure and functional state of AtAKT1 in a lipid bilayer. **a)** Molecular model of the AtAKT1 homotetramer with a single subunit colored by domain. **b)** Sharpened electrostatic potential map, colored by domain as in **a** with low-pass filtered envelope shown to depict the nanodisc and poorly-resolved Ankyrin repeats. **c)** Thede-de-active voltage sensing domain with charge transfer center residues shown. **d)** Pore profile of the closed channel with pore-lining residues on the P-loop and S6 shown including the selectivity filter near the extracellular side, and activation gate at the intracellular side. The pore radius is shown plotted with respect to the distance along the channel axis, calculated using MOLE ^25^. **e)** Atomic model of the selectivity filter and K^+^ ion superpositions, with sharpened electrostatic potential map shown zoned 2.5 Å around the atoms of interest. **f)** Residues that form the selectivity filter from AtAKT1 and other K^+^ channels, highlighting the signature GYG motif.

The overall architecture of the AKT1 channel is similar to other non domain-swapped ion channels such as the hyperpolarization-activated HCN1 ^12,13^, KAT1 ^14^ and LliK ^15^ channels, the depolarization-activated ether-a-go-go channel ^16^, and ligand-gated CNGA1 channel ^17^. In these channels, the S4 helix from the voltage sensing domain (VSD) that bears the gating charges is linked to the S5 helix of the pore domain via a short loop (**1a**) such that the pore and voltage sensing domains of the same subunit coalesce together as a bundle. This is in contrast to the canonical domain swapped architecture found in *Shaker*-like, Na_V_, and two-pore channels, in which the VSD and pore helices are separated by a long helical S4-S5 linker ^13^. Our structure is consistent with a de-active, closed channel in which the gating charges on S4 are in a resting, ‘up’ position (**1c**), and the intracellular activation gate is constricted to preclude the diffusion of ions through the central pore (**1d**). Previous work on the hyperpolarization-activated HCN1 channel showed that two gating charges are transferred downwards across the hydrophobic constriction site (HCS) during activation, accompanying a helical break in S4 ^13^. In our structure, the corresponding charges are R164 and R167, which rest above the HCS formed by F100, while R169 and R170 are below the HCS closely interacting with an intracellular negative cluster formed by E61, D103 and D139 (**1c**).

The selectivity filter of AKT1 is nearly identical to all other K^+^-selective channels of known structure, consisting of a tetrameric “bracelet” arrangement of backbone carbonyls from the signature TVGYG motif (**1f**), mimicking the 8 water ligand field around K^+^ ions in bulk solvent ^18,19^ (**1d,e**). We assign four K^+^ ions to densities on the channel axis within the selectivity filter which correspond to super-positions of alternately populated pairs of sites (**1e**). In addition, we structured a crown of four extracellular water molecules partially hydrating a S_0_ K^+^, which are possibly hydrogen bonded to histidine residues (H260), conserved in plant inwardly-rectifying K^+^ channels (**S5a,b**) ^20^, that line the extracellular face of the channel. In this model, H260 donates a hydrogen bond from its δ nitrogen to its own backbone carbonyl, poising it’s putatively un-protonated ε nitrogen towards a crown water proton, thus aligning the water towards the S_0_ K^+^ (**S5a,b**). The H260 imidazoles may facilitate partial dehydration of incoming K^+^•(H_2_O)_8_ complexes by positioning the hydration crown, allowing the K^+^ to transition into a partially hydrated state (i.e. K^+^•(H_2_O)_4_(C=O)_4_) above the selectivity filter. Below the selectivity filter is an aqueous bath where K^+^ ions can re-hydrate, followed by an intracellular activation gate wherein the sidechains of I285 and T289 form an ion-excluding constriction point (**1d**).

Initial image processing with imposition of C4 symmetry showed high resolution features in the transmembrane domain, including an abundance of annular phospholipids, K^+^ ions in the selectivity filter and pore waters, but smeared density corresponding to intracellular domains. For this reason, we re-performed all 3D image processing without imposition of symmetry and also four-fold symmetry expanded the consensus C4 particle stack. Subsequent 3D classification produced multiple conformations of the channel, each exhibiting a dramatic reduction of symmetry from four to two-fold. Intriguingly, we observe the C-linker, which links the pore domain to the CNBHD, in two distinct configurations on the same C2-symmetric reconstruction. Neighboring AKT1 subunits adopt alternating C-linker conformations: one conformation is termed ‘flat’, similar in conformation to that seen in LliK ^15^ and HCN1 ^12^, while the second conformation is termed ‘kinked’ and resembles that of KAT1 ^14^. Variations in Clinker and CNBD / CNBHD architecture have been observed between different channels though the exact nature of these differences is not always clear ^7,12,14,15,17,21^. Our structures suggest that these channels may sample multiple conformations during their respective activation cycles, and that there may be direct implications for regulation of electrical activation (**2a,b,c,d**).

**Figure 2.**
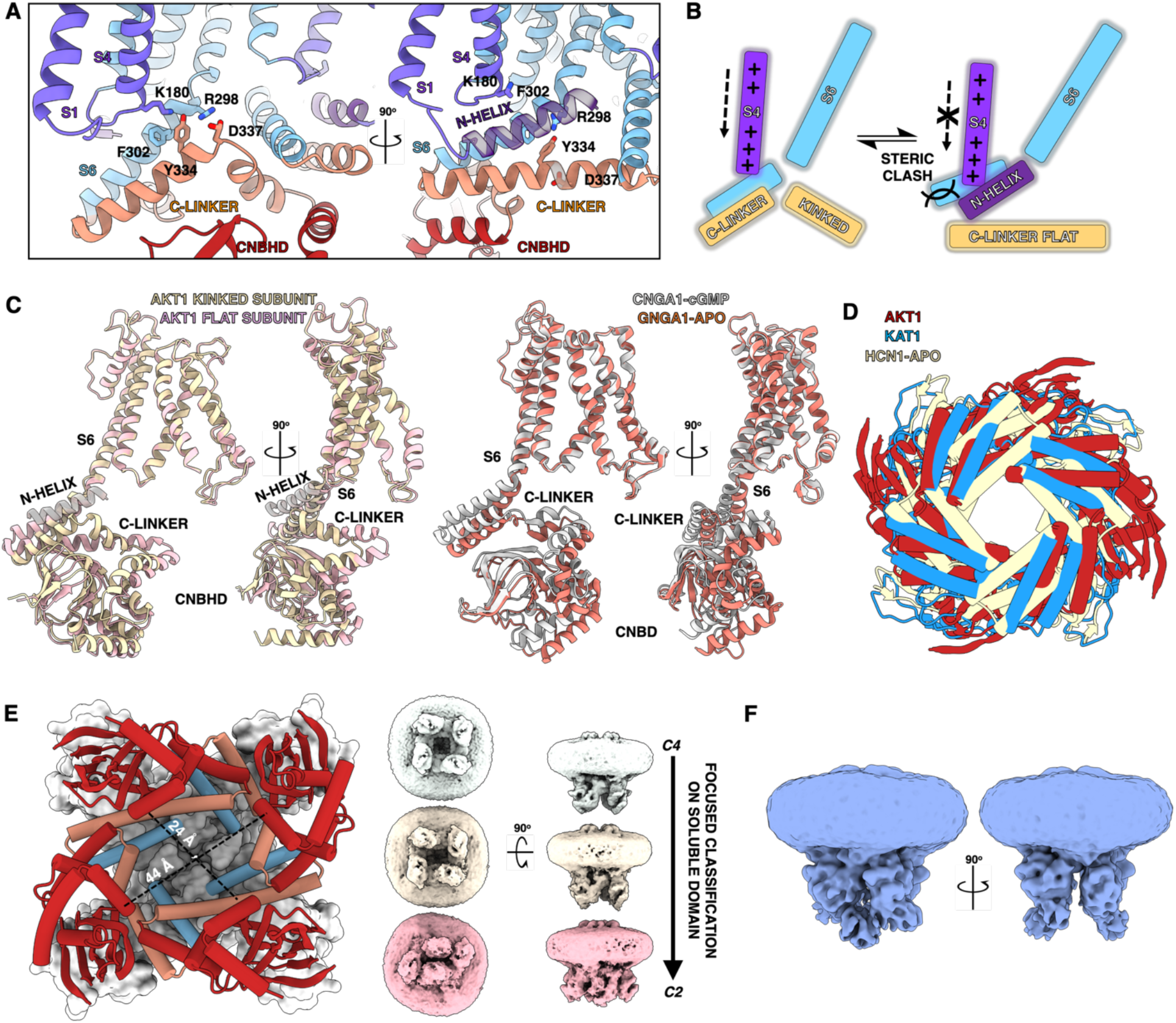
Symmetry reduction in AtAKT1 and its implications for voltage activation. **a)** The C-linker adopts two different conformations in the C2-symmetric structure. In one conformation, the linker is kinked, forming a nexus with S4 and S6. In the adjacent subunit, the C-linker is flattened by the insertion of the pre-VSD N-terminal helix, breaking most atomic interactions in the nexus. **b)** Graphical model of how N-helix insertion flattens the C-linker (represented by solid arrows) and potentially interferes with the downward translocation of S4 during membrane hyperpolarization (represented by dashed arrows). **c)** Orthogonal views of overlaid AKT1 kinked and flat subunits (left), and apo- and holo-CNGA1 (PDBs: 7LFT and 7LFW, respectively) highlighting differences in C-linker and CNBD conformations. **d)** Overlay of the CNBD/CNBHDs of AKT1, KAT1 (PDB: 6V1X), and apo-HCN1 (PDB: 5U6O), viewed along the channel axis from the intracellular side. **e)** Three conformations of the CNBHD domains are recovered from 3D classification (right), the C4 class displayed at top, mid-C2 class center, and the ultra-C2 class at bottom, all viewed at low contour. The soluble region in the mid-C2 class was resolved to high resolution and was used to build an atomic model of AKT1, shown on the left. The degree of asymmetry is quantified by the distances between D337 on opposite C-linkers. **f)** Orthogonal views of the mid-C2 map, displayed at low contour to show the Ankyrin domains in two distinct configurations: one pair exhibits extensive inter-domain interactions while the neighboring pair does not touch.

In the kinked conformation, the C-linker forms an intricate interaction nexus with the VSD and S6 (**2a**). D337 is situated at the summit of the kink and forms an apparent salt bridge with R298 on S6. Y334 forms H-bonds with K180 at the bottom of S4 and R298, and also an aryl stacking interaction with F302 on S6. In the flat conformation, this nexus is significantly altered due to the intercalation of an N-terminal helix that precedes S1 (**2a,b**). This N-helix is only observed above the flat C-linker, where it makes extensive contacts with S6, the VSD, and the C-linker itself. The residues connecting this helix to S1 are poorly resolved, precluding accurate register assignment and atomic-level description of how helix insertion flattens the C-linker.

The N-helix is notably excluded from the kinked C-linker sites, indicating that it is in a flexible, unresolved conformation. In the flat conformer, the aryl interaction between Y334 and F302 is largely broken, as is the salt bridge between D338 and R298 as D337 is forced to face away from S6 (**2a,e**). Although the exact functional consequence of these differences is still unclear, we propose that the insertion of the N-helix serves to autoregulate the electrical activation of S4. In order to activate the channel, S4 must translocate downwards to some extent ^13,14,22^, and the presence of the N-helix between it and the C-linker may sterically occlude this process (**2b**).

Asymmetry is further observed in subsequent domains. The CNBHD that follows the flat C-linker is notably flexible, whereas the post-kinked CNBHD is resolved to high resolution allowing for accurate atomic structure determination. As a result of this asymmetry, the inter-CNBHD interactions around the channel ring differ considerably. Focused classification on the CNBHD domains from the original C1 particle stack of EM images without imposition of symmetry recovered three distinct states. The first of these has apparent C4 symmetry while the latter two have varying degrees of C2 symmetry (**2e**). At low contour levels, the Ankyrin domains are also seen to adopt a strongly-elongated C2 symmetric configuration in which two adjacent pairs of Ankyrin domains form close contacts with one another leaving a gap between these and the alternate pair of domains (**2f**). As the channel was heterologously expressed in mammalian cells without a CIPK/CBL10 phosphorylation system, we assume that the Ankyrin domains are unmodified. Therefore, it’s feasible that Ankyrin phosphorylation alters this configuration for channel activation, possibly by breaking apart these inter-Ankyrin interactions to symmetrize the channel.

Although the precise role of symmetry breaking in a degenerate ligand-binding domain remains unclear, our results suggest that it might serve a regulatory role. The observed differences between adjacent CNBHD conformations is reminiscent of the transition between apo- and cNMP-bound states of cyclic nucleotide-gated ion channels (**2c,d**), posing the possibility that this domain in AKT1 has been converted from a ligand-binding domain to a site of allosteric communication between the Ankyrins and the channel’s pore. It is enticing to compare our observations with those of other CNBHD-containing channels whose electrical properties are altered by N- and C-terminal regulatory domains. In the case of hERG, an N-terminal helix acts in concert with the CNBHD to enhance the channel’s activation kinetics and mar its speed of deactivation ^23,24^. In the depolarization-activated hERG channel, insertion of an N-terminal helix between its C-linker and VSD would, according to our model, ‘push’ the S4 helix upwards towards an activated conformation, and prevent its deactivation into a resting, downwards state. Inversely, downwards translocation of S4 during activation in a hyperpolarization-activated channel such as AKT1 would be inhibited by the presence of such a helix. We hope that the unusual findings presented in this manuscript will help inform experiments on this very important plant channel, and on the many salient hyperpolarization- and CNBD-containing-channels in human physiology.

## Acknowledgements

We thank Phuong Nguyen for assistance with mammalian cell culture. We thank David Bulkley, Glen Gilbert and Zanlin Yu for their support of the UCSF electron microscopy core, and the NIH grants that support them. We thank Paul Thomas, Matt Harrington, Joshua Baker-LePain and the Wynton HPC team for computational assistance. We thank Janet Finer-Moore, Evan Green and Richard Aldrich for critical reading of the manuscript. M.S.D. acknowledges an NSF graduate research fellowship. Research was supported by NIH grant GM24485 (to R.M.S.).

## References

1. Hedrich, R. Ion channels in plants. Physiol. Rev. 92, 1777–1811 (2012).

2. Ward, J. M., Mäser, P. & Schroeder, J. I. Plant ion channels: gene families, physiology, and functional genomics analyses. Annu. Rev. Physiol. 71, 59–82 (2009).

3. Sharma, T., Dreyer, I. & Riedelsberger, J. The role of K(+) channels in uptake and redistribution of potassium in the model plant Arabidopsis thaliana. Front. Plant Sci. 4, 224 (2013).

4. Hirsch, R. E., Lewis, B. D., Spalding, E. P. & Sussman, M. R. A role for the AKT1 potassium channel in plant nutrition. Science 280, 918–921 (1998).

5. Li, J. et al. The Os-AKT1 channel is critical for K+ uptake in rice roots and is modulated by the rice CBL1-CIPK23 complex. Plant Cell 26, 3387–3402 (2014).

6. Anderson, J. A., Huprikar, S. S., Kochian, L. V., Lucas, W. J. & Gaber, R. F. Functional expression of a probable Arabidopsis thaliana potassium channel in Saccharomyces cerevisiae. Proc. Natl. Acad. Sci. USA 89, 3736–3740 (1992).

7. Matulef, K. & Zagotta, W. N. Cyclic nucleotide-gated ion channels. Annu. Rev. Cell Dev. Biol. 19, 23–44 (2003).

8. Lee, S. C. et al. A protein phosphorylation/dephosphorylation network regulates a plant potassium channel. Proc. Natl. Acad. Sci. USA 104, 15959–15964 (2007).

9. Hedrich, R. & Kudla, J. Calcium signaling networks channel plant K+ uptake. Cell 125, 1221–1223 (2006).

10. Xu, J. et al. A protein kinase, interacting with two calcineurin B-like proteins, regulates K+ transporter AKT1 in Arabidopsis. Cell 125, 1347–1360 (2006).

11. Ren, X.-L. et al. Calcineurin B-like protein CBL10 directly interacts with AKT1 and modulates K+ homeostasis in Arabidopsis. Plant J. 74, 258–266 (2013).

12. Lee, C.-H. & MacKinnon, R. Structures of the Human HCN1 Hyperpolarization-Activated Channel. Cell 168, 111–120.e11 (2017).

13. Lee, C.-H. & MacKinnon, R. Voltage Sensor Movements during Hyperpolarization in the HCN Channel. Cell 179, 1582–1589.e7 (2019).

14. Clark, M. D., Contreras, G. F., Shen, R. & Perozo, E. Electromechanical coupling in the hyperpolarization-activated K+ channel KAT1. Nature 583, 145–149 (2020).

15. James, Z. M. et al. CryoEM structure of a prokaryotic cyclic nucleotide-gated ion channel. Proc. Natl. Acad. Sci. USA 114, 4430–4435 (2017).

16. Whicher, J. R. & MacKinnon, R. Structure of the voltage-gated K^+^ channel Eag1 reveals an alternative voltage sensing mechanism. Science 353, 664–669 (2016).

17. Xue, J., Han, Y., Zeng, W., Wang, Y. & Jiang, Y. Structural mechanisms of gating and selectivity of human rod CNGA1 channel. Neuron 109, 1302–1313.e4 (2021).

18. Zhou, Y., Morais-Cabral, J. H., Kaufman, A. & MacKinnon, R. Chemistry of ion coordination and hydration revealed by a K+ channel-Fab complex at 2.0 A resolution. Nature 414, 43–48 (2001).

19. Morais, J. H. Energetic optimization of ion conduction rate by the K+ selectivity filter. Cabral

20. Hoth, S. et al. Molecular basis of plant-specific acid activation of K+ uptake channels. Proc. Natl. Acad. Sci. USA 94, 4806–4810 (1997).

21. Li, M. et al. Structure of a eukaryotic cyclic-nucleotide-gated channel. Nature 542, 60–65 (2017).

22. Dai, G., Aman, T. K., DiMaio, F. & Zagotta, W. N. The HCN channel voltage sensor undergoes a large downward motion during hyperpolarization. Nat. Struct. Mol. Biol. 26, 686–694 (2019).

23. Muskett, F. W. et al. Mechanistic insight into human ether-à-go-go-related gene (hERG) K+ channel deactivation gating from the solution structure of the EAG domain. J. Biol. Chem. 286, 6184–6191 (2011).

24. Gustina, A. S. & Trudeau, M. C. hERG potassium channel gating is mediated by N- and C-terminal region interactions. J. Gen. Physiol. 137, 315–325 (2011).

25. Pravda, L. et al. MOLEonline: a web-based tool for analyzing channels, tunnels and pores (2018 update). Nucleic Acids Res. 46, W368–W373 (2018).

